# Identifying and tracking mobile elements in evolving compost communities yields insights into the nanobiome

**DOI:** 10.1101/2023.02.02.526783

**Authors:** Bram van Dijk, Andrew D. Farr, Pauline Buffard, Franz Giersdorf, Jeroen Meijer, Bas E. Dutilh, Paul B. Rainey

## Abstract

Microbial evolution is driven by rapid changes in gene content mediated by horizontal gene transfer (HGT). While mobile genetic elements (MGEs) are important drivers of gene flux, the nanobiome – the zoo of Darwinian replicators that depend on microbial hosts – remains poorly characterised. New experimental approaches and analyses are necessary to advance our understanding beyond simple pairwise MGE-host interactions. To detect horizontal transfer, a bioinformatic pipeline (xenoseq) was developed to cross-compare metagenomic samples, which was then applied to metagenomic data from evolving compost communities. These communities were routinely exposed to an “MGE cocktail” derived from allopatric communities. We show that this results in horizontal transfer of a multitude of previously undetected MGEs, including bacteriophages, phage-plasmids, megaplasmids, and even nanobacteria. Sequences that spread from one community to another are shown to disproportionally carry characteristics of phages and insertion-sequences, *i*.*e*., traits of canonically parasitic MGEs. We also show that one particularly prolific mobile element - a 313 kb plasmid – correlates substantially with rates of ammonia production, which under nitrogen limitation is likely beneficial. Taken together, our data show that new experimental strategies combined with bioinformatic analyses of metagenomic data stand to provide insight into the drivers of microbial community evolution.

## Introduction

Horizontal gene transfer (HGT) can markedly affect the evolutionary fate of microbes[1]– [3]. Besides transformation — where bacteria directly take up environmental DNA — all horizontal movement of genetic material is catalysed by mobile genetic elements (MGEs), Darwinian entities with dynamics of their own[4]. Over the last century, a whole zoo of MGEs has been observed, ranging from the distinctly parasitic phages[5]–[9] and transposons[10]–[12], to MGEs like plasmids[13]–[15], integrative and conjugative elements (ICEs)[16]–[19], and repetitive extragenic palindromic doublets forming hairpins (REPINs).[20], [21]

The relationships between MGEs and their hosts are complex, ever changing, and highly context-dependent. For example, while conjugative elements are typically benign, they may also promote their own survival at the expense of hosts[19], [22]. Similarly, while bacteriophages are typically predatory or parasitic, they can be co-opted to benefit hosts[5], [23]–[25]. Moreover, various mobile elements have been shown to recombine, and even parasitise other mobile elements[26]–[30]. Clearly, MGEs have highly fluid compositions, roles, and eco-evolutionary consequences, mediating a constant flux of DNA through microbial communities. This flux of DNA has been compared to community-level sex[31], [32]. Taken together, these processes may result in collections of locally adapted genes, rather than locally adapted microbial species[33], [34]. Understanding this process requires a multi-level perspective that takes into account not just microbiomes, but also *nanobiomes*: all the biologically relevant entities much smaller than bacteria.

New methods are essential to shine light on the “dark matter”[35]–[37] of nanobiomes. Traditional microbiology allows for the identification of novel MGEs when they are linked to ecologically relevant traits. For example, ICEs were discovered by observing nodulation of a non-native legume in New Zealand fields by indigenous rhizobia[38], [39, p. 500]. ICEs have also been identified by observing the movement of antimicrobial resistance[40], [41] and heavy metal resistance genes[18]. However, such routes to discovery depend on both abilities to culture focal microbes and carriage of selectable phenotypic traits.

Recently, the discovery of MGEs has been fuelled by metagenomics through culture-independent assembly of DNA replicons without prior assumptions about biological relevance. Moreover, bioinformatic tools have been developed that can separate a wide range of possible MGEs from microbial hosts[42]–[49]. Although such tools allow differentiation of chromosomal DNA from phages, plasmids, and other MGEs, discovery of MGEs is constrained by existing databases and training sets that are based on known MGEs. Moreover, metagenomic detection of MGEs *per se* does not directly yield insights into the ecological relevance of the element. In summary, unbiased detection and characterisation of unknown MGEs remains a major challenge.

Recently, Quistad *et al*. reported an experimental approach to investigate the impact of MGEs on the ecology and functioning of complex microbial communities[50]. In the experiment, MGE cocktails were routinely collected from independent compost communities, pooled, and then redistributed across mesocosms (see **Figure 1a**). This simple manipulation resulted in MGEs being repeatedly exposed to new hosts thus enhancing their evolutionary potential and facilitating horizontal dissemination of ecologically relevant genes. The affected communities were labelled “horizontal” and contrasted with a paired set of “vertical” communities that received no MGE cocktail. Communities were then subjected to metagenomic sequencing, which - by virtue of the experimental design - allowed the identification of sequences in the horizontal communities that were not detected in the respective ancestral communities (hereafter referred to as “unique sequences”), and therefore likely represent MGEs derived from allopatric mesocosms. While many of these unique elements were shown to be phage-like, their detection was not dependent on matches to existing databases, but rather, appearance among the set of “unique sequences”. However, as pointed out in the original study, many “unique sequences” did not show features typical of phages or other known MGEs, leaving open the possibility that these candidate sequences had no history of horizontal dissemination. Instead, these sequences may have been rare elements that were selected for simply due to changes in species composition. For example, sequences may become detectable (and thus classified as unique), solely through an increase in frequency placing them above the threshold for metagenomic detection. While the study by Quistad *et al*. marked a significant conceptual advance in evolutionary thinking regarding microbial communities, much remained unexplored with respect to the identity of unique sequences.

**Figure 1:**
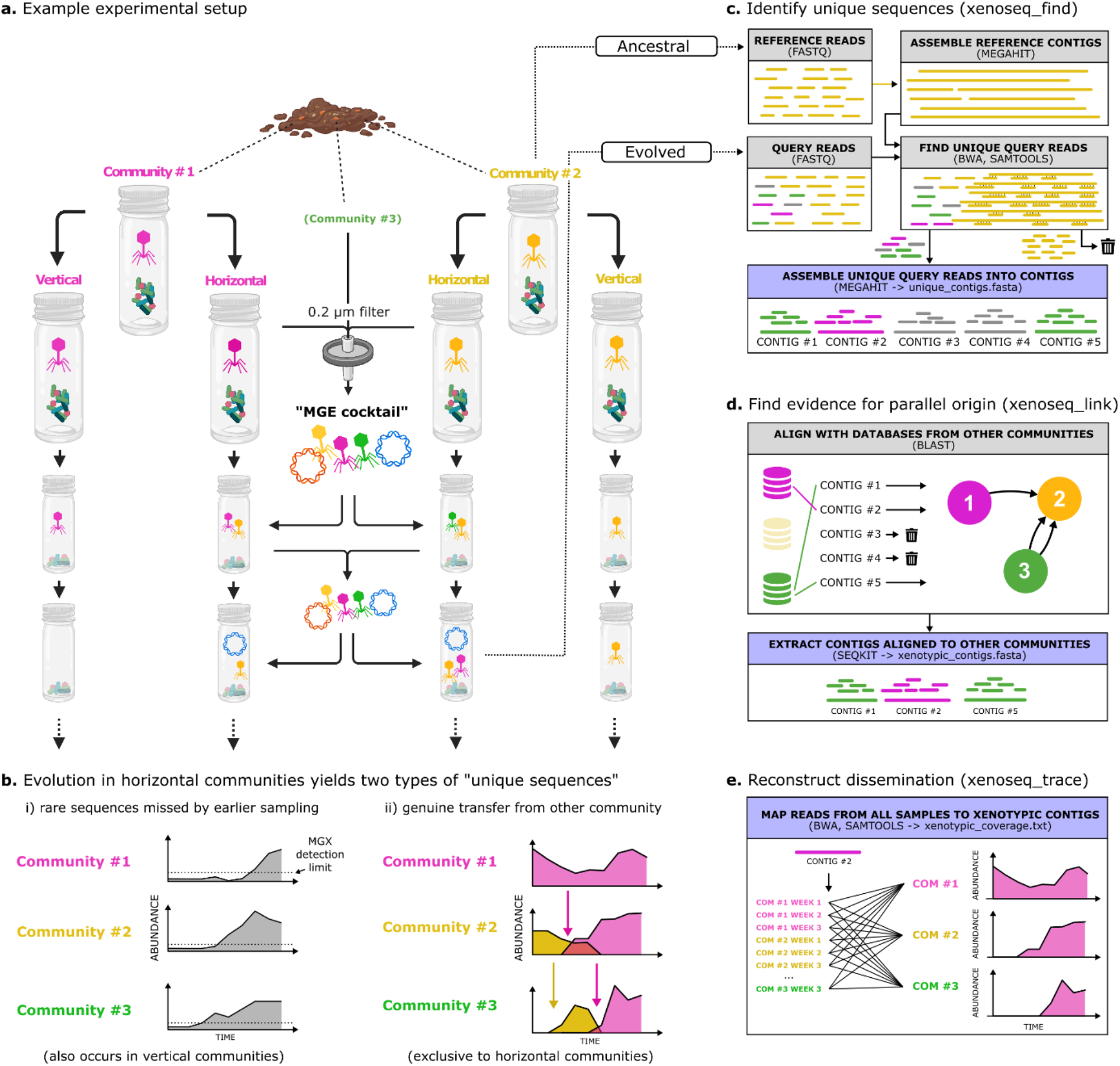
Overview of a) experimental setup, b) sources of unique sequences, and c-e) the bioinformatic pipeline. a) Experimental evolution protocol to identify novel MGEs and other mobile sequences (Quistad et al, 2020). Multiple parallel communities were established from garden compost, which were then propagated on cellulose as sole carbon source and serially transferred every two weeks. Horizontal treatments were provided with an “MGE cocktail” which was derived by pooling material collected from samples passaged through 0.2 μm filters. This manipulation allows MGEs and linked genes to transfer horizontally from one community to the next. b) Sequencing samples obtained from mesocosms can yield two types of “unique sequences”. Shown are cartoons to visualise how some unique sequences (left) are the result of rare sequences that were undetected by earlier sampling, while other unique sequences (right) are the result of genuine transfer from another community. c) The first subroutine of xenoseq (xenoseq*_*find) compares raw DNA sequences reads in fastq format from one sample (query, hereafter called *evolved*) with another sample (subject, hereafter *ancestral*). Ancestral samples are assembled into contigs, which are used as bait to remove reads from the evolved samples. Unmapped reads from the evolved samples are then assembled into “unique contigs”, stretches of DNA that newly appeared in the community. d) To distinguish between two sources of unique sequence shown in panel b, the second subroutine (xenoseq_link) identifies promising candidate mobile elements by aligning them to all other ancestral communities. The subset of contigs that align well to allopatric communities are referred to as *xenotypic contigs*. e) Finally, the dissemination patterns of xenotypic contigs is reconstructed by mapping DNA sequences reads from all communities against the contig. Sequencing depth and breadth are stored in a tabular text file.

Here, we describe a broadly applicable pipeline (github.com/bramvandijk88/xenoseq) that takes as input, raw sequencing reads from time series experiments, and detects newly introduced sequences. When applied to metagenomic data from Quistad et al.’s experimental protocol described above, xenoseq detects movement of a large variety of candidate MGEs. One important advance is the ability to distinguish between sequences that are selected due to demographic changes in patterns of species abundance, from those that genuinely appear to be horizontally disseminated from an allopatric community. The latter are hereafter referred to as “xenotypic sequences”. We show that xenotypic sequences are enriched in recognisable components of phages and IS-elements, that is, canonical selfish genetic elements. We also observe the transfer of a 313 kb mobile element with plasmid-like features, whose abundance correlates with changes in ammonia production rates. Finally, by re-analysing data from the Quistad *et al*. experiment, we show that horizontal communities show repeated shifts in important cellulose degrader ecotypes, especially favouring a nitrogen-fixing strain of the genus *Cellvibrio* that appears to be the host of the aforementioned 313kb mobile element detected by xenoseq. Our work confirms that opening up communities to the movement of MGEs has profound eco-evolutionary effects, and contributes a new method to identify candidate MGEs in a relatively unbiased way.

## Materials and Methods

### Bioinformatic pipeline to detect transfer events

The xenoseq pipeline (https://github.com/bramvandijk88/xenoseq, **Figure 1c-e**) is a wrapper that combines read trimming, assembly, read mapping, read filtering, and local alignment to seek evidence of horizontal transfer between communities. This pipeline takes as input raw (untrimmed) fastq files containing shotgun metagenomic data of derived samples (query, in which to look for newly introduced sequences) and datasets from ancestral samples (reference). Query reads are trimmed using fastp^55^ (v0.23.2), mapped against the corresponding reference contigs using BWA[51], [52] (v0.7.17) using default options, and samtools[53] (v1.15.1) is used with the flag “*-4”* to extract unmapped reads. These reads are assembled into “unique contigs” using Megahit[54] (v1.2.9) (*xenoseq_find*, **Figure 1c**). Next, these unique contigs are blasted against a local database using NCBI Blast[55] (v.2.13.0) of all other reference communities to link the emergence of unique contigs to allopatric “donor” communities. Linked contigs are extracted from the unique contigs using Seqkit[56] (v.2.3.0) (*xenoseq_link*, **Figure 1d**). By default, contigs are linked to a donor when blast has at least one high-scoring segment with a minimum length of 300 with at least 99% nucleotide identity. While the algorithm and cut-offs limit the sensitivity of the algorithm at larger evolutionary distances, they are designed to detect sequences that have recently diverged and are still largely identical. The contigs that match target sequences in an allopatric community are referred to as “*xenotypic contigs*”. Finally, to detect the shifts in abundance of the detected contigs, reads from all query and reference samples are mapped (xenoseq_trace), and coverage/breadth statistics are saved in a tab-separated file. This allows for visualisation of transfer across communities (**Figure 1e**). In this manuscript, we further analyse sequences >10kb in length.

By default, xenoseq runs all subroutines (*xenoseq_find, xenoseq_link, xenoseq_trace*), but each subroutine can be run independently by modifying the command-line flags. Finally, xenoseq uses GNU parallel^59^ to run multiple jobs simultaneously.

### Xenoseq benchmark

To benchmark the pipeline, the introduction of MGEs into mock-communities was simulated (see **Supplementary Material I** full details). Six bacterial genome sequences were downloaded from RefSeq. Then, two mock-communities were generated, one with an even taxon distribution (easy dataset), and one with a highly skewed taxon distribution (hard dataset). Simulated MGE sequences were either i) randomly inserted into genome sequences, representing integrative elements, or ii) included as separate replicons linked to a single host genome by adding them to the genome fasta file as a separate contig. Illumina sequencing was simulated from the resulting “communities” with ART^62^, using both default and ten-fold elevated error rates, which were used as input to benchmark the pipeline. Simulation of mock-communities and horizontal gene transfer events was done in R using the packages biostrings^60^ and seqinr^61^. Another (small) mock-community dataset is available on the repository, and can be used to rapidly test whether the pipeline and its dependencies are configured correctly.

### Metagenome-assembled genomes

To obtain genome sequences of important players in the communities, we generated Metagenomic-Assembled Genomes (MAGs). By combining samples across multiple time points, we improved the potential detection of rare types whose coverage in a single sample is insufficient for assembly. Quality control of the sequencing reads was done with Prinseq version 0.20.4 [57] using *“–derep 14 -lc method -c threshold 20”*. We trimmed adapters with Flexbar v.3.5.0[58] using the “*–adapter-preset Nextera -ap ON*” flag. Combined reads from multiple time points for each community were cross-assembled *de novo* using metaSPAdes v.3.14.0[59]. Reads from each sample were mapped back to the assembled contigs using bwa-mem v.0.7.17-r1188[51], and coverage calculations were performed with SAMtools v1.7[53]. Contigs were binned using metaBAT2 v2.12.1 [60] and MAG quality was assessed with CheckM v1.1.3[61]. All MAGs and contigs were annotated using BAT and CAT (Bin/Contig Annotation Tool, respectively, v.5.1.2)[62], which uses prodigal[63] to predict open reading frames and diamond[64] to query open reading frames against the NR database [65]. When BAT or CAT could not reliably assign a taxon due to conflicting annotations between ORFs, we assigned a higher order taxonomic rank to the MAG: that is, to genus level if species identity could not be assigned, family level if genus could not be assigned, *etc*. CAT was called using parameters *“--index_chunk 1 --block_size 5 --top 11”* using the CAT database and taxonomy constructed on 2021-04-30. Read mapping of samples from individual time points (above) was used to study the relative abundance of individual MAGs.

### DNA Degradation assay

To address the question as to whether the identified sequences are derived from living cells or DNA liberated by lysed cells, we tested the ability of microbial compost communities to degrade extracellular DNA. To do this, four experimental microbial communities were established from samples of four independent compost heaps. Compost was washed in M9 salt solution and stored with glycerol saline at -80 °C. Frozen stocks were thawed on ice and 4.25 mL of the stock was then washed of glycerol by two successive cycles of centrifugation (4000 g, ten minutes) and resuspension (in five mL sterile M9 salt solution), followed by a final resuspension in 1mL M9 salt solution. The washed cells were then added to 100 mL bottles with 19 mL M9 media and a piece of cellulose paper as a carbon source. Mesocosms were incubated without shaking at 28 °C for 14 days to allow community growth. The entire volume including any remaining paper was then transferred to 50 mL centrifuge tubes, vortexed for five minutes to produce a slurry, and 1 mL of slurry was transferred to fresh M9 media for another 14 days of incubation.

During this time a stock of genomic DNA (gDNA) from *Escherichia coli* (REL606) from ∼72 mL of overnight culture was made using the “DNeasy Ultraclean Microbial kit” (Qiagen), resulting in a stock of ∼30 mL of gDNA at 20 μg/mL (stored at -20 °C). REL606, nor any other *E. coli* strains occur naturally in the compost, so tracking this gDNA will allow the persistence of extracellular DNA to be quantified below.

After 14 days of culture, the entire volume of each composting culture was vortexed to a slurry, and one mL of slurry was then added to 19 mL of M9 medium to initiate four replicate mesocosms. Unlike previous transfers, each 20mL culture was spiked with ten μg of *E. coli* REL606 gDNA. Three community-free controls were established with ten μg gDNA spiked into nine replicate 20 mL tubes containing only M9 medium. For time point t=0, the contents of one mesocosm and one control was immediately transferred to a 20 mL centrifuge tube following spiking, reduced to a slurry and then centrifuged at 4000 g for five minutes. The supernatant was transferred to 15 mL sterile tubes and frozen at -80 °C for later DNA extraction. This process was repeated after 24 hrs, 48 hrs and 14 days, with four replicate communities and three community-free controls at each time-point.

Prior to extraction of DNA from the samples, cultures of *Pseudomonas fluorescens* SBW25 were made and DNA extracted (using the same kit as for *E. coli*) providing a stock of 50 ng/μL (again, kept frozen at -20 °C). Frozen samples of the communities were thawed on ice, and a 3mL sample was spiked with 30 μL of *P. fluorescens* gDNA. At t=0, when no degradation of DNA has occurred, samples contain approximately comparable concentrations of *E. coli* and *P. fluorescens* gDNA. Because *P. fluorescens* gDNA is added daily, its concentration should be constant, while *E. coli* gDNA is expected to be degraded by the community.

To determine rates of DNA degradation and consumption by communities, spiked samples were vortexed and samples were filtered using a 0.2 μm syringe-filter. DNA of 2 mL of filtrate was isolated using Phage DNA Isolation Kit (Norgen Biotek Corp) by following the manufacturer’s instructions, except for loading two (rather than one) mL sample through a single purification column to improve DNA yield. Extracted DNA was then sequenced using a NextSeq MidOutput 300 cycle run resulting in 2x150 bp paired-end reads **(see Supplementary Table I)**. Reads were aligned to the non-redundant (NR) protein database[65] with Diamond^54^ using the blastx subroutine, where the top hit was used to identify the read as being either *P. fluorescens, E. coli*, or “other” (the latter representing either species present in the communities or erroneously assigned reads).

### MGE classification

We identified 1756 candidate MGEs in horizontal communities (>10kb, linked to at least one allopatric “donor” community) and analysed with a variety of tools for predicting different kinds of MGEs. Viral sequences were predicted using Vibrant (v1.2.0)[42], Virsorter2 (v2.2.3)[43], Seeker (v1.0.3)[44], and CheckV (v0.8.1)[66]. While CheckV is primarily designed to estimate completeness of candidate viral genomes, we found that sequences with either low, medium or high scores were typically identified as viral by other tools as well (see **Figure 3** in main text). Plasmid sequences were predicted using PlasFlow (v1.1.0)w[45] and PlasClass (v0.1)[46] and, if relevant, the origin of replication with OriFinder 2022[67]. ICEs were identified with ICEfinder[47]. IS elements were predicted with ISEscan (v1.7.2.3)[49]. Integrons were predicted using Integronfinder2 (v2.0rc6)[48]. All open reading frames on MGEs were annotated with prokka[68] with both the default database and the PHROG[69] database for identification of viral genes. All tools were run with default options, results are shown in ***Supplementary Table 1, Sheet 3***.

**Figure 2:**
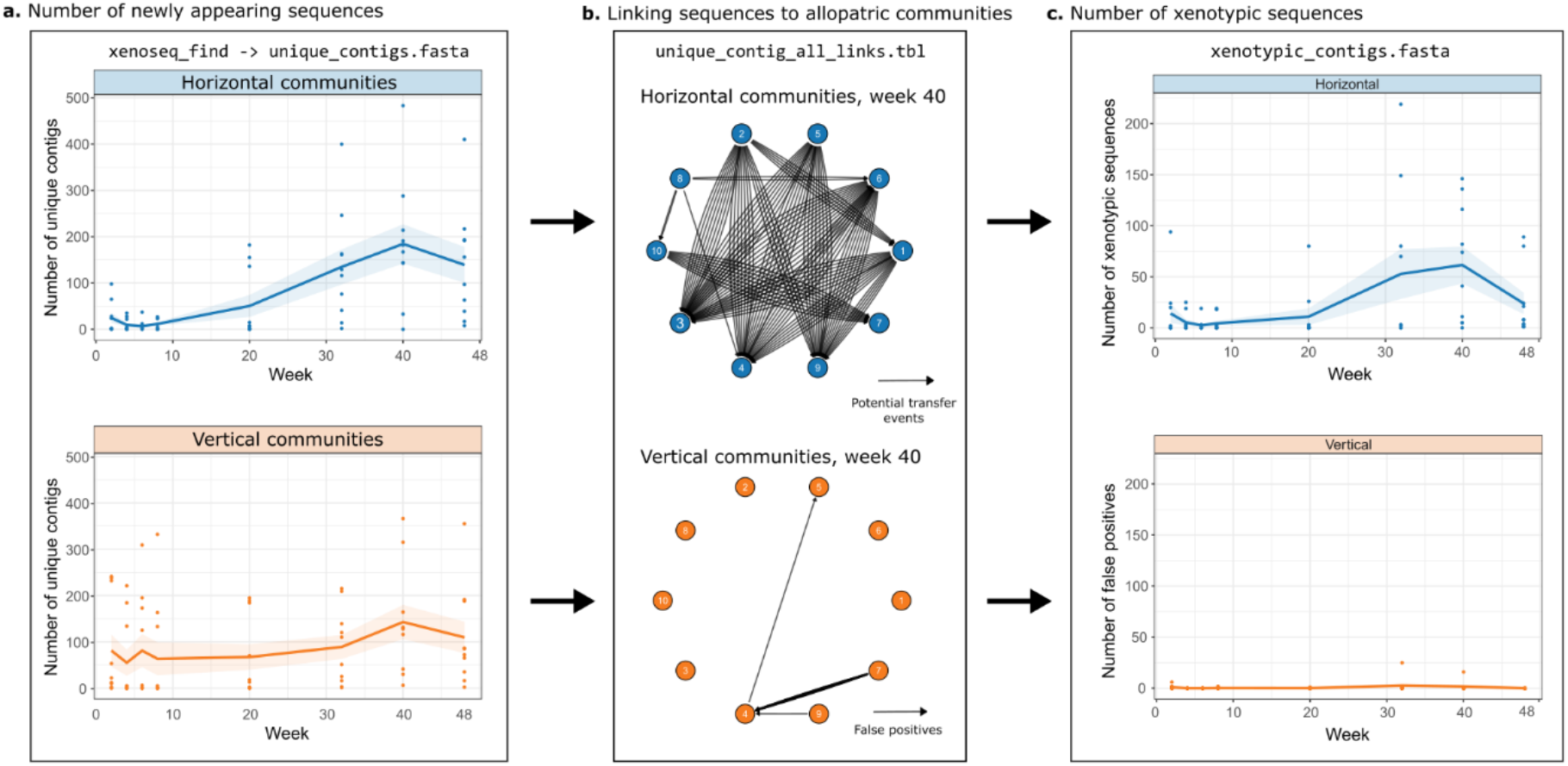
Horizontal communities are enriched in xenotypic sequences. a. Newly appearing contigs within horizontal (blue) and orange (orange) communities identified by xenoseq_find. The solid lines represent the mean across 10 communities, and the shaded area represents the standard error. b. Aligning contigs identified by xenoseq_find to ancestral samples allows for linking of donor-acceptor pairs (example network shown for horizontal and vertical communities at week 40, also see **Supplementary Figure 1+2**). For horizontal communities, links in the network represent elements that are allopatrically selected and transferred. For vertical communities, the links instead represent false positives, as transfer from allopatric communities is excluded by the experimental design. c. The number of xenotypic contigs (sequences that could be linked to at least one donor community) identified in horizontal communities is shown in blue, and false positives observed in vertical communities is shown in orange. Dots represent 10 different replicate communities, the solid lines represent the mean a cross communities, and the shaded area represents the standard error.

**Figure 3:**
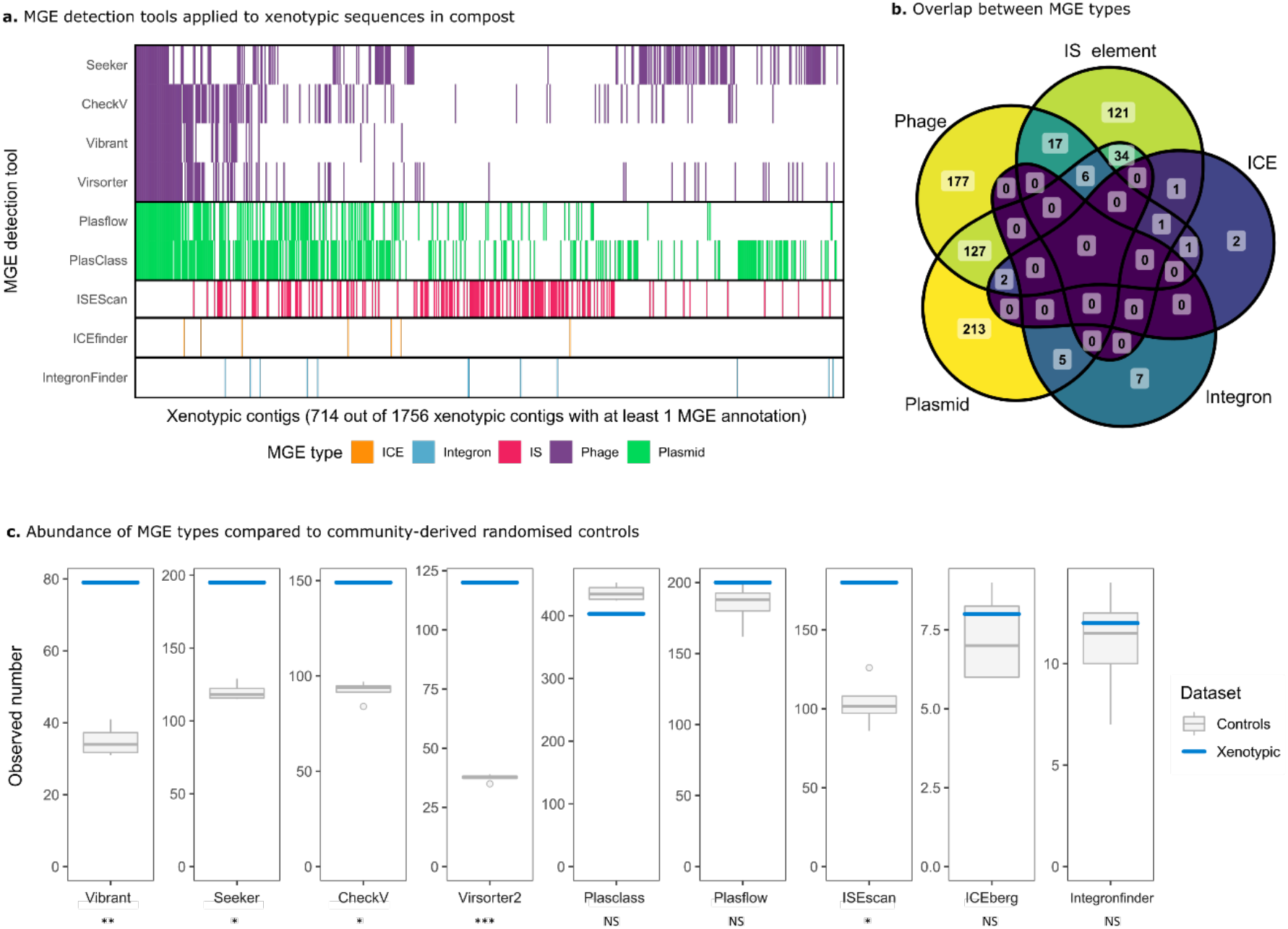
MGE detection tools reveal a diverse set of mobile elements in compost communities. a. Various tools were used to predict MGEs including phages, plasmids, IS-elements, ICEs, and integrons in xenotypic sequences. On the x-axis are 714/1756 candidate MGE contigs with at least one MGE prediction. On the y-axis are the MGE tools. A coloured bar indicates that the contig was predicted by the corresponding tool. b. A venn diagram shows that there is substantial overlap between MGEs, especially phages and plasmids. This may indicate interactions or hybridisation of phages and plasmids, but could also be the result of erroneous assignment by one of the tools. c. The number of MGE annotations derived from xenotypic sequences is compared with four controls. Controls consist of contigs with similar lenghts from the xenotypic contigs, but are sampled from the whole compost community metagenome. A modified T-test (Crawford-Howell[77]) was used to infer whether phages, plasmids, IS-elements, ICEs, or Integrons were over- or under-represented in the xenotypic sequences (* = p-value < 0.05, ** = p-value < 0.01, *** = p-value < 0.001).

### MAG metabolic functions

The limiting resources in the M9 medium of the experiment by Quistad *et al*. (2020) are carbon (provided by the cellulose paper) and nitrogen (1 mM ammonium chloride from M9 salts), both of which are added every 14 days during serial passaging. To get insight in the community function of lineages, we screened each MAG for their ability to metabolise these two resources. Although many relevant proteins are involved, we focus attention on processes that contribute most to the community-level function in compost, such as endoglucanases and cellobiosidases (degrading cellulose), nitrogenases (nitrogen fixation), and nitrate/nitrite reductase (nitrogen reduction). More “private” functions, such as the ability to take up glucose, glycolysis, and downstream pathways, are not taken into consideration for this study. Protein sequences were extracted from the Uniprot database (keywords: “*endoglucanase*”, “*cellobiosidase*”, “*nitrate reductase*”, “*nitrite reductase*”, and “*nitrogenase*”, all with the additional query “*reviewed:true*”) and aligned to the MAGs using diamond blastx[64] with default settings. A score was assigned to each MAG by calculating the fraction of queries hit. For example, when one out of two cellobiosidases from Uniprot gave a significant hit, a score of 0.5 was assigned. All scores are given in **Supplementary Table 1, Sheet 5**. For **Figure 5b**, a metabolic function was assigned when the score was greater than 0.1. When two MAGs with the same CAT annotation occurred in multiple communities, but the metabolic scores were not identical due to differences in genomic coverage, the highest score was used.

**Figure 4.**
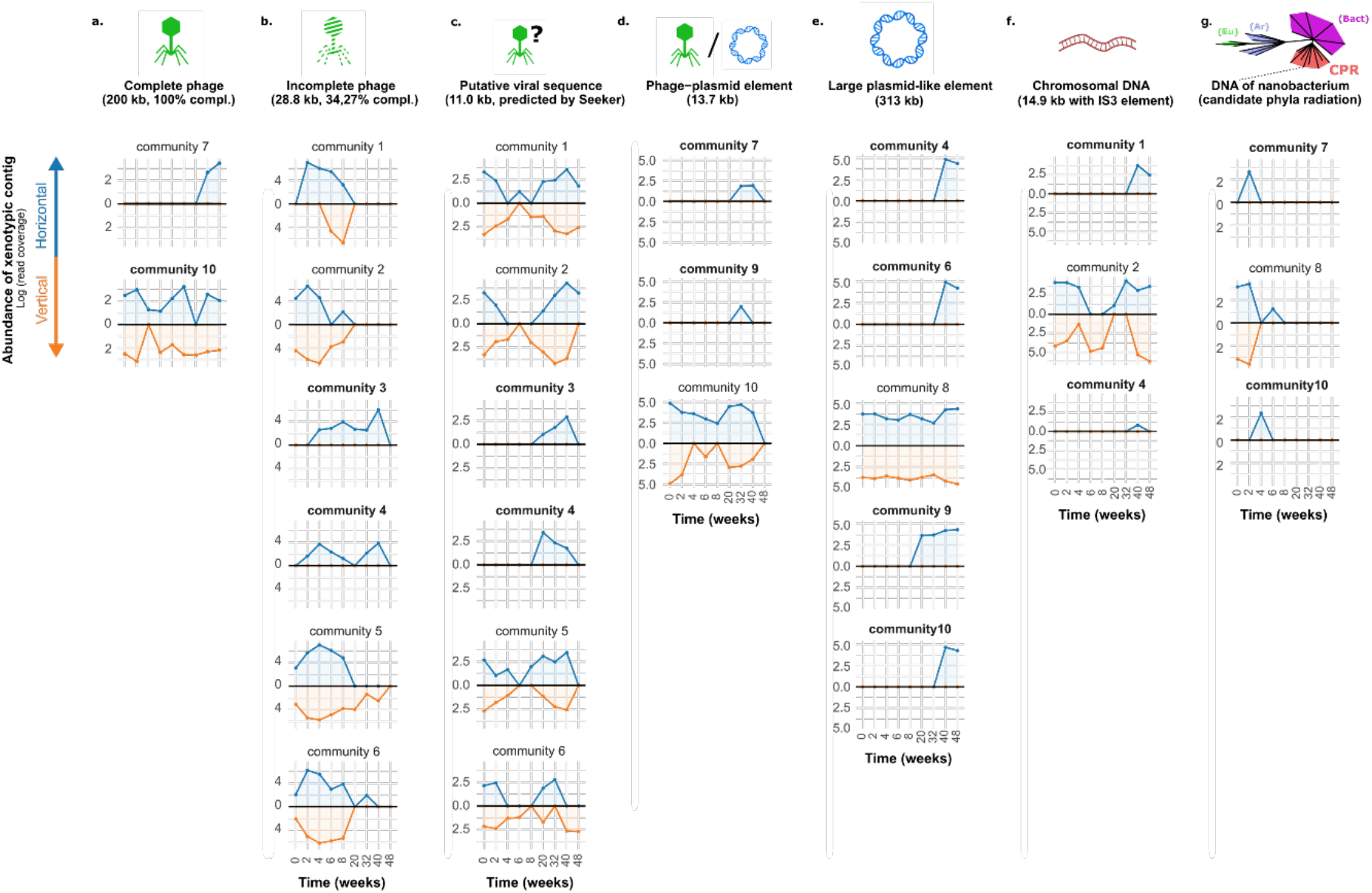
Tracing selected xenotypic sequences across communities. Results of xenoseq_trace for seven xenotypic sequences; namely (**a**) a phage with 100% completion as reported by CheckV (abbreviated as compl), (**b**) an incomplete yet highly abundant phage sequence (at certain time points covered by > 10^*6*^ reads), (**c**) a putative viral sequence exclusively predicted by seeker, (**d**) a sequence reported to be both phage and plasmid by all relevant MGE tools, (**e**) a large plasmid that successfully establishes in four communities, (**f**) a putatively chromosomal region flanked by a transposable element, and (**g**) an sequence not predicted to be an MGE, annotated by CAT as being *Candidatus saccharibacterium*. For all panels, abundances (y axes, log scaled read coverage, mean per nucleotide) are shown across communities over all time points (x axes). Abundance in horizontal communities is shown in blue, whereas abundance in vertical communities is shown on the opposing axes in in orange. Communities in which the sequence is unique to the horizontal regime are shown in bold. Communities in which the contig was not observed are omitted for clarity. The full interactive data set is available as Supplementary Material.

**Figure 5.**
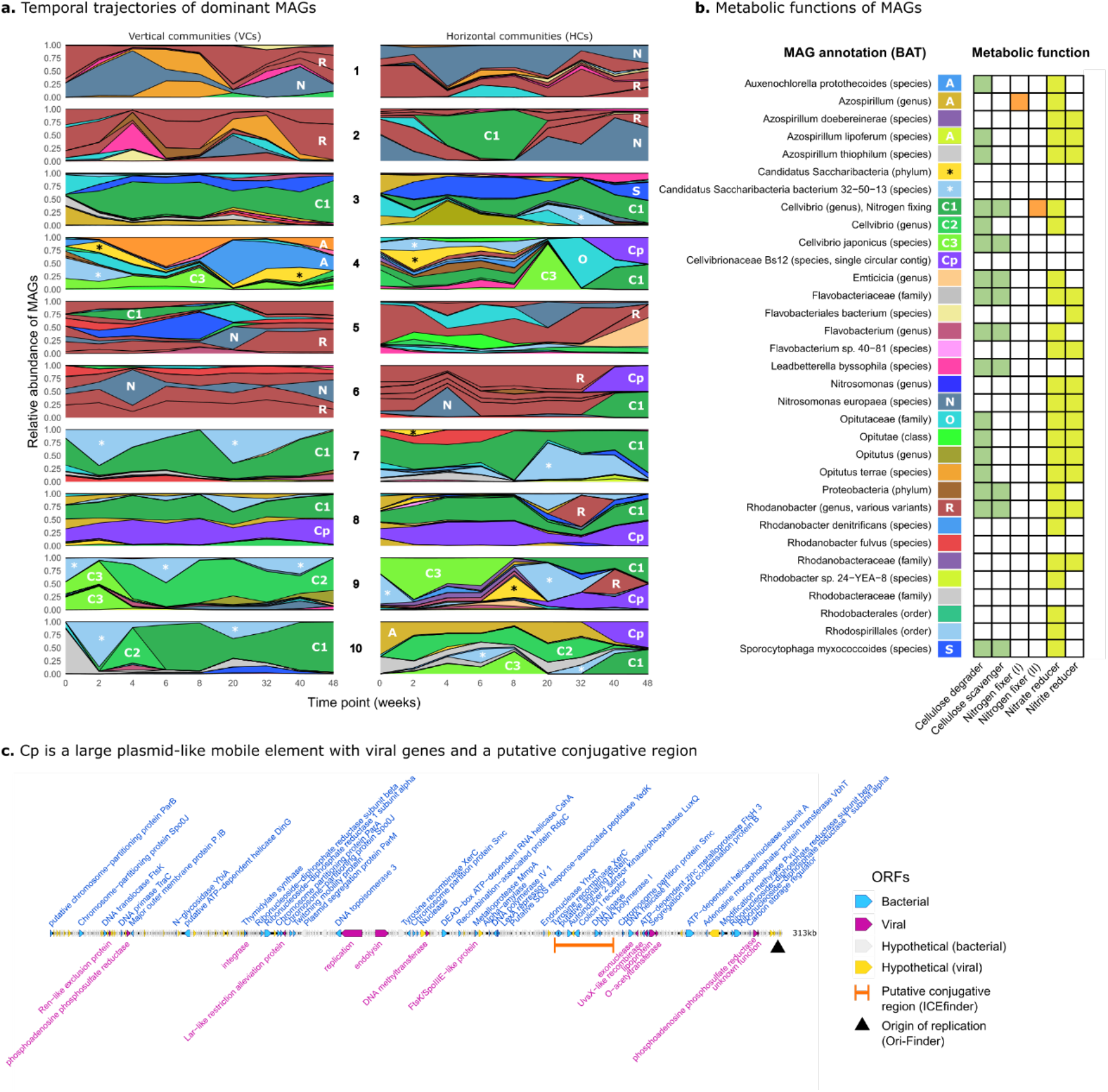
Horizontal communities show repeated shifts dominant MAGs. a. For all 20 replicate experiments (10 horizontal, 20 vertical), abundance (relative read coverage) is shown for MAGs over time. For clarity, only MAGs that are highly abundant across multiple time points are shown (representing on average 59.6% of the total community), and abundance is given as the fraction of reads mapping only to these particular MAGs. A subset of MAGs are labelled with a letter (C1, R, N, Cp, etc.) corresponding to the legend in (**b**) for guidance. b. Metabolic functions assigned to each MAG. MAGs with evidence for encoding endoglucanases, cellobiosidases, nitrogenases (I and II), and nitrate/nitrite reductase were assigned roles to cellulose degraders, cellulose scavengers, nitrogen fixers, nitrate reducers, and nitrite reducers, respectively (see Methods). A score was calculated based on the number of significant hits to these protein sequences, and MAGs with a score greater than 0.1 are coloured in the heatmap. c. A circular, 313kb mobile element (Cp) that emerges in four independent horizontal communities. The element contains both plasmid-like and phage-like traits.

### Measuring MGE cocktail DNA concentration

To measure the DNA concentration present in DNA cocktails, compost mesocosms were established using the protocol described in Quistad *et al*. (2020). Briefly, samples were obtained from a compost heap (Plön, Germany) in September 2021. Five grams from each compost sample was transferred into a 100 ml glass flask containing a four cm^2^ piece of cellulose paper as a complex carbon source (Whatman cellulose filter paper) in 20 mL minimal M9 medium which contains 0.935 mM ammonium chloride. Subsequently, the founder mesocosms were incubated without shaking at 28°C for two weeks. The lids remained slightly opened, allowing gas exchange.

After a two-week incubation period, the cellulose paper was transferred into a falcon tube with 20 ml of minimal M9 medium and vortexed into a slurry. The slurries were vortexed, and 12 ml of cellulose slurry was centrifuged, for 10 minutes, at 4000 g. Subsequently, 10 ml supernatant was filtered through a 0.2 μm filter to produce the MGE cocktail. For DNA extraction, MGE cocktails were concentrated using an ultracentrifuge at ∼26500 g for 45 minutes. The supernatant was discarded and the pellet resuspended in ∼2 mL medium. DNA was extracted using the Phage DNA Isolation kit (Norgen Biotek Corp) or QIAprep Spin Miniprep Kit, following the manufacturer’s protocol. The amount of extracted DNA was assessed by Qubit HS assay. The measurements are found in **Supplementary Table I, Sheet 4**.

### Data analysis and visualisation

Data analysis and visualisation was done in R[70], using the packages ggplot2[71, p. 2], dplyr[72], ggplotly[73], patchwork[74], gggenomes[75], ggraph (github.com/thomasp85/ggraph), and rtracklayer[76].

## Results

We developed a bioinformatic pipeline that aims to identify novel MGEs and other biologically relevant entities from metagenomic data. The pipeline does not rely on existing databases, but instead detects transfer of candidate MGEs based on sequences newly introduced into evolving communities. As such sequences are of “foreign” nature, we refer to these as “xenotypic sequences”, and the pipeline is called xenoseq (https://github.com/bramvandijk88/xenoseq). For a full description of the bioinformatic pipeline see methods and **Supplementary Material I and II**.

Xenoseq was first benchmarked using simulated mock communities (***Supplementary Materials II*)**, and then applied to datasets from Quistad *et al*. (2020). The data by Quistad *et al*. is of particular interest, because the experimental design allows for dissemination of MGEs within *horizontal communities*, without allowing the migration of microbial cells (**Figure 1a**). As short-read metagenome samples were prepared from all communities at various timepoints, it is possible to identify sequences that are not native to the evolved communities, that is, candidates for transfer of phages, plasmids, or other MGEs. However, not all sequences that newly appear are due to horizontal transfer between allopatric communities. False positives can emerge when a sequence grows in abundance sympatrically, which can happen with rare species that are initially below the metagenomic detection limit (**Figure 1b**). This false positive signal due to sympatric forces ought to apply equally to *horizontal* and *vertical* communities. Hence, after identifying newly emerged “unique sequences” (xenoseq_find, **Figure 1c**), further evidence for transfer is sought by aligning these contigs against sequences from allopatric communities (xenoseq_link, **Figure 1d**). Then, as a final step, read mapping provides further insight into the origin and dissemination of xenotypic contigs (see **Figure 1e**).

Note that xenoseq can in principle be applied to datasets that deviate from the particular experimental design by Quistad *et al*., including natural samples. The only requirement is that longitudinal samples are taken from parallel communities which undergo exchange of MGEs or other biologically relevant entities.

### Horizontal compost mesocosms are enriched in xenotypic sequences

The mesocosms from the study by Quistad *et al*. (2020) represent a “challenging case” for the xenoseq pipeline, because of the unprecedented diversity and many rare types. As above-mentioned, shifts in abundance of rare types (as observed by Quistad *et al*., 2020) can generate false positives (**Figure 1b**). As a control, we therefore also run xenoseq for the vertical communities, which are “closed” vertical communities. Indeed, we found that unique sequences emerge in both horizontal and vertical communities (**Figure 2a**, a total of 5617 and 6883 sequences respectively). However, only horizontal communities contained unique contigs that could be linked to allopatric communities (**Figure 2b**). In total, 1756 (31.2%) of contigs in horizontal communities could be linked to an allopatric “donor” community, versus only 58 (0.8%) of contigs from vertical communities. Thus, all evolved compost communities show sequences that are amplified over time, but evidence for allopatric origins of these sequences is only found in horizontal.

A possible confounding factor in the detection of xenotypic sequences could arise if DNA included in the MGE cocktail were to remain intact during the 14-day incubation period, is sequenced, and ends up in the metagenomic data. To determine whether such extraneous DNA is problematic, we firstly determined the concentration of DNA in MGE cocktails that were prepared following the protocol described in Quistad *et al*. (2020). At DNA concentrations of 0.67 ng/ml in the MGE cocktail, we consider it highly unlikely that such sequences appear in the metagenome 14 days later, unless they are further amplified during that period. Secondly, we tested whether added genomic DNA remained detectable after introduction into compost mesocosms and incubation for 14 days. In the absence of communities, DNA remained intact and detectable at 14 days, however, genomic DNA added to communities was undetectable 48 hours after addition (**Supplementary Figure 3**). These data demonstrate that, in order for xenoseq to detect xenotypic contigs, their DNA sequences must be substantially amplified within the recipient community. Such amplification indicates some form of selection for the candidate MGE, *e*.*g*. because it is a selfish element or conveys a substantial fitness benefit to a host.

### Xenotypic sequences are enriched for phages and IS-elements

The MGE cocktail that was distributed among mesocosms could contain bacteriophages, plasmids, naked DNA, membrane vesicles, and potential other (unknown) vehicles of transfer. We subjected our 1756 xenotypic sequences to a variety of MGE detection tools, which provided predictions as to the sequences being viral[42]–[44], [66], plasmid^45,46^, IS-carrying^49^, ICE-carrying^47^, or integron-carrying^48^. We found that 714 (40.6%) of xenotypic sequences received at least one MGE prediction (see **Figure 3A**). Considerable overlap was found between phages and plasmids (**Figure 3B**), but also between the other elements. These results indicate that there is substantial recombination between MGEs, but also highlight the challenges of unambiguously predicting the MGE type from sequence by using the selected tools.

To test whether xenotypic sequences are enriched in MGEs, these data were compared to an arbitrary set of sequences with a similar length distribution, sampled from the metagenomic contigs. Phages and IS-elements appear to be over-represented on xenotypic sequences, whereas the numbers of plasmids, ICEs, and integrons are not significantly different (**Figure 3C** and **Supplementary Table I, Sheet 3**). As only canonically selfish elements (phages, IS-elements) are enriched, these results further corroborate the importance of sequence amplification after introduction into a new community.

Interestingly, after applying nine state-of-the-art tools for predicting five categories of MGEs, 1042 xenotypic contigs (59.4%) remained unidentified. These sequences could constitute the transfer of chromosomal DNA or unknown MGEs, or alternatively, could be erroneously inferred to be transferred horizontally.

### Movement of xenotypic sequences between communities

To investigate the dissemination of xenotypic contigs, xenoseq maps reads from all communities to the xenotypic contigs representing candidate MGEs to track their dynamics. **Figure 4** shows the results for seven selected examples: a complete 200 kb phage (**Figure 4a**), an incomplete but prolific phage sequence (**Figure 4b**), a putative viral sequence exclusively predicted by Seeker (**Figure 4c**), an element predicted as both phage and plasmid unanimously (**Figure 4d**), a large 313kb plasmid (**Figure 4e**), and a stretch of apparent chromosomal DNA adjacent to an IS3 element (**Figure 4f**). Finally, xenoseq also revealed transfer of chromosomal DNA fragments lacking MGE predictions, many of which were annotated as Candidatus *Saccharibacterium* (**Figure 4g**), and many other interesting sequences (full interactive dataset available online).

### Compost mesocosms form distinct community types

Thus far, we have illustrated that treating evolving microbial communities with an “MGE cocktail” derived from all communities promotes the movement of various MGEs, and even nanobacteria between communities. We have also shown that xenotypic contigs are especially enriched in SGEs, suggesting the importance of independent sequence amplification for the survival of MGEs after their introduction into allopatric communities. In the following sections, we examine the ecological and evolutionary consequences of this treatment. To investigate whether and how the horizontal communities are distinct from the vertical communities, we studied the abundance of Metagenome-Assembled Genomes (MAGs). These MAGs were assigned a taxonomic classification using the Bin Annotation Tool (BAT^52^). When taxonomic rank could not be determined reliably due to conflicting ORFs, a higher-order taxonomic rank was assigned instead, ensuring all MAGs had a classification.

Each MAG was screened for various genes related to two metabolically relevant functions: cellulose degradation and nitrogen metabolism (see Methods and **Supplementary Table I, Sheet 5**). The relative abundance of dominant MAGs is shown for each community in **Figure 5a** (for an interactive graph of all MAGs, see **Supplementary Files**). The MAGs are dominated by either *Rhodanobacter* (shown in brown in **Figure 5a**) or *Cellvibrio* lineages (shown in green). Members of both these genera are able to degrade cellulose, the sole carbon source in the mesocosms (**Figure 5b**). As many other MAGs do not have this ability, *Rhodanobacter* and *Cellvibrio* appear to occupy a similar niche as the primary degrader, which could explain why they appear to be mutually exclusive. The two community types furthermore have distinct lineages that coinhabit the mesocosms. For example, the communities dominated by *Rhodanobacter* lineages often coexist with *Nitrosomonas europaeae* (steel blue), which can reduce nitrate and nitrite (**Figure 5b**). The communities dominated by *Cellvibrio* instead frequently host species of C. *Saccharibacteria* (denoted by an asterisk in **Figure 5a-b**), a nanobacterium from the candidate phyla radiation[36] (CPR). The ∼800 kb genome of this nanobacterium shows no capacity to degrade cellulose or reduce nitrogen sources (**Figure 5b**), indicating dependency on other species in the community. Co-occurrence analysis (see **Supplementary Figure 4**) suggests that this nanobacterium is either an epiphyte or parasite of *Cellvibrio*, with massive blooms hinting at the latter (see **Supplementary Figure 4b**).

### MGE cocktails can disrupt established degrading types

As can be seen from **Figure 5a**, vertical communities (VCs, left) are relatively stable in the composition of dominant MAGs, establishing either with *Cellvibrio* or *Rhodanobacter* as the primary cellulose degrader. Horizontal communities (HCs) however show rapid shifts in these primary degraders. For example, HC2 initially establishes with *Rhodanobacter* (brown) as the primary degrader. However, during week four to eight, a *Cellvibrio* lineage (C1, green) becomes more dominant relative to other MAGs. Finally, R*hodanobacter* re-emerges as the primary degrader. *Vice versa*, HC8 and HC9 are initially established as *Cellvibrio*-communities, and show a transient appearance of a *Rhodanobacter* lineage after 32/40 weeks.

While the above-mentioned disruptions caused by the MGE cocktails are transient, *Cellvibrio* C1 in HC6 appears to completely overturn the ecosystem structure that was established for during the preceding 32 weeks. *Cellvibrio* C1 also emerges in three other mesocosms independently (in HC9 at week 8, and HC4 and HC10 at week 32), and its emergence is always accompanied by a MAG containing only a single 313kb contig (shown in purple in **Figure 5a**, henceforth called Cp). This contig is of circular nature, and is in fact the same plasmid-like element earlier identified by xenoseq (**Figure 4e**). The large mobile element carries important plasmid partitioning proteins(*e*.*g*. parB and parM) typically associated with low copy-number plasmids[78], [79] (see **Figure 5c**), which is consistent with it co-occurring with *Cellvibrio C1* in an approximate 1:1 ratio. In addition to features that are akin to large plasmids, Cp also carries many ORFs associated with phages as evident in matches to the viral PHROGs-database^66^ (for example, integrase and endolysin). Finally, Cp carries a putative conjugative region identified by ICEfinder. As this plasmid is present in earlier time points only in VC/HC8, this suggests this community as the donor. We hypothesise that Cp transfers horizontally, enabling Cellvibrio C1 (the putative host) to displace the previously established ecosystem structure.

### Cp+ communities accumulate more ammonia

Because cellulose is the only exogenously provided carbon source in the mesocosms, the ability to degrade cellulose is found in many MAGs (see **Figure 5b**). While all the *Cellvibrio* lineages have the ability to degrade cellulose, *Cellvibrio* C1 is unique in also carrying a nitrogen fixing enzyme (flavoprotein, EC 1.19.6.1). The ability to fix nitrogen is likely important, because 1 mM ammonium chloride from the M9 medium added every 14 days is the only other source of nitrogen. Indeed, by plotting the abundance of *Cellvibrio* C1 with previously published data on ammonia production (Quistad *et al*., 2020), we found a strong positive correlation (**Figure 6a**). Interestingly, this positive correlation was weaker - or at times entirely absent - in VCs, where the Cp plasmid was absent in all but one community. Similarly, when focussing on HCs where Cp was absent (HC3 and HC7), no positive correlation was observed (**Supplementary Figure 4**). These data suggest that Cp promotes the fixation of nitrogen in *Cellvibrio C1*.

**Figure 6.**
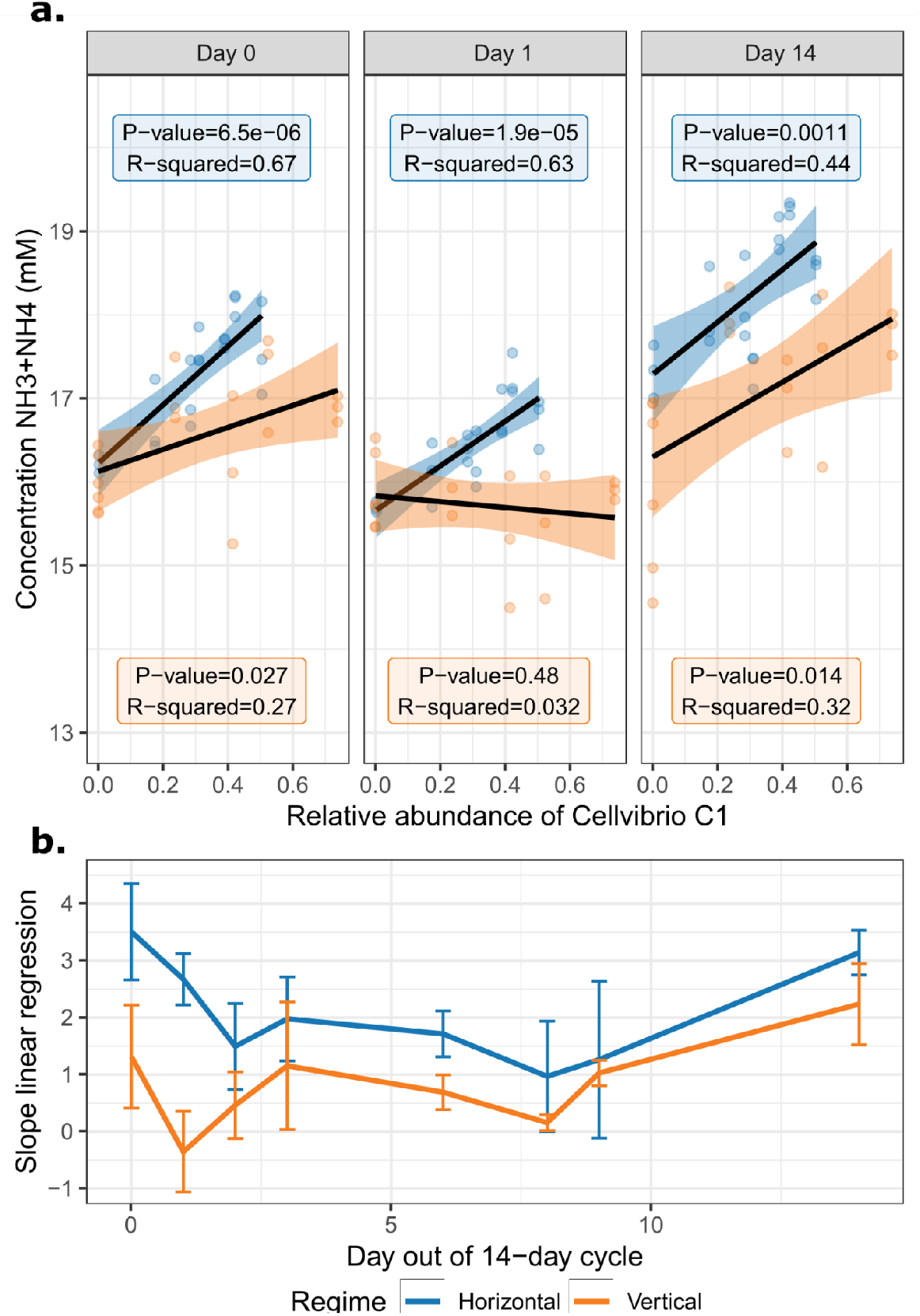
*Cellvibrio* C1 fixes more nitrogen in Cp+ (horizontal) communities. a. The relative abundance of *Cellvibrio* C1 was determined by the proportion of reads from the sample at week 48 mapping against this MAG, which was then plotted against ammonia production measurements from Quistad *et al*. (2020). *Cellvibrio* C1 was present in four vertical communities and seven horizontal communities, yielding a total of seven abundance values for *Cellvibrio C1*, plotted on the x-axis. For each of these values, three technical replicate measurements of ammonia were performed at several time points in the 14-day cycle. The y-axis shows the measured ammonia concentrations (NH3+NH4). A linear regression model was fitted to the data points. For day 0, 1 and 14, this fit is shown with their corresponding p-values and R^*2*^ values (see **Supplementary Figure 5** for all days). b. The slope of the linear regression is plotted for all days in the bottom panel, revealing that horizontal communities consistently show a steeper slope, suggesting that the large Cp plasmid promotes nitrogen-fixation by the *Cellvibiro* C1 MAG. Error bars denote the standard deviation of the slope across three technical replicates of ammonia production measurements.

## Discussion

Mobile genetic elements (MGEs) are important for microbial ecology, evolution, and community function. Moreover, as diverse as microbiomes are, the nanobiome - all the Darwinian replicators dependent on microbial hosts – may be even more diverse. We developed a bioinformatic pipeline called xenoseq to shine light on the nanobiome, inspired by experiments from Quistad et al. (2020). Using time-resolved metagenomic data, xenoseq can distinguish between two sources of novel sequences, (i) those that arise due to local demographic changes, and (ii) those that are the consequence of horizontal transfer between parallel communities. We show that the latter category, called *xenotypic sequences*, require amplification after introduction into allopatric communities, which may be enabled by independent replication as parasitic selfish elements or as beneficial elements that are linked to a particular host. We found that xenotypic sequences were especially enriched in selfish genetic elements (phages and IS elements). While other MGEs such as plasmids and ICEs were not enriched, we found a 313 kb plasmid that appears to convey a substantial fitness benefit to its host. Taken together, our data shows that the pipeline can successfully identify a broad range of interesting MGEs, without any prior knowledge as to their identity.

The experimental strategy of Quistad et al. (2020) and our bioinformatic pipeline can be applied to any microbial community and can accommodate specific selection pressures, such as antimicrobial resistance[80], [81], heavy metal resistance[82], or bioremediation capacity[83]. Furthermore, even without any evident selection pressures, the method can provide direct evidence of selection pressures experienced by communities, by investigating which traits are transferred. Quistad et al. observed the enrichment of nitrogen metabolism genes in horizontal communities, which were here able to link to the proliferation of a large 313 kb plasmid-like sequence. Given its apparent indirect role in fixing nitrogen, we hypothesise that the observed amplification of the large element confers a fitness advantage (**Figure 6**), but questions remain as to the mechanism of transfer. We argue that, although the possibility cannot be excluded, it seems unlikely for such a large element to transfer as naked DNA. An alternative route of transmission for large plasmids could be packaging inside membrane vesicles (MVs)[84]–[87], as was recently shown in *Klebsiella pneumoniae*[88].

Besides the movement of MGEs themselves, our study also suggests competition between ecologically relevant species is enhanced in horizontal communities. This enhanced competition may be due to “kill the winner” dynamics[89], where phages preferentially bring down successfully established microbial species to improve their own evolutionary fate, making niche space available for competitors. A similar dynamic was observed in a recent study that illustrates how the induction of prophages can significantly impact the assembly of chitin-degrading communities[90]. Gaining a further understanding of these effects may help us establish healthy microbial communities in societally relevant systems such as the gut[91] or plant rhizosphere[92], [93]. Taken together, understanding how MGEs influence the competition between microbial species and the robustness of microbial communities (or lack thereof)[94]–[96], may enable improved evolutionary predictions[97].

An unexpected finding of our study was the transmission of a CPR nanobacterium of the phylum *Saccharibacteria* (previously known as TM7). Based on microscopy or genomic analysis, other CPR bacteria have been suggested to be epibionts, such as *Vampirococcus lugossi*, which appears to attach to photosynthetic bacteria[98]. Indeed, close relatives of this MAG are not much larger than a phage^75^, and the reduced genome (∼800 kb) is indicative of a symbiotic relationship with another species. Our co-occurrence analysis (see **Supplementary Figure 4a)** revealed a potential connection with a bacterial lineage belonging to the genus *Cellvibrio*. Rather than an epibiont, we suggest that this TM7 member may also be an intracellular parasite, as the boom-and-bust dynamics shown and long periods of stasis shown in **Supplementary Figure 4b** are reminiscent of viral dynamics. The identification of this nanobacterium, and the newly generated hypothesis of a host-parasite relationship, highlight an additional power to our approach. Excitingly, *Cellvibrio* can be isolated and cultured, paving the way for future studies on this nanobacterium and the interactions with its host.

Other annotation-free methods to track HGT in microbial ecosystems have recently been developed, relying for example on differential read coverage[99], assembly graphs[100], [101], discordant read pair mapping[102], or Hi-C metagenomics[80], [103], [104]. These new approaches allow detecting the transfer of genomic regions, while our xenoseq pipeline and the long-term community evolution experiments aim to detect the transfer of candidate MGEs between communities. Thus, compared to our approach, earlier methods do not provide insights into the dynamics and functional consequences of MGE transmission. Our strategy combines experimental intervention, metagenomics and bioinformatic analysis to discover elements capable of transmission as nanoparticles and subsequent amplification, without prior knowledge of the elements themselves. In other words, using our method, it becomes possible to interrogate the nanobiome free from pre-conceived notions of the identity of its members.

### Concluding remarks

Our work confirms earlier conclusions that microbial ecosystems are greatly influenced by the flux of DNA created by MGEs, even when these fluxes are initially subtle. Hence, we argue that understanding the nanobiome – the zoo of Darwinian entities much smaller than bacteria – is crucial to understanding microbial ecology, and how these systems eventually scale up to impact entire ecosystems.

## Data availability

Raw sequencing data are taken from Quistad *et al*. (2020), and are publicly available online (https://www.mg-rast.org/mgmain.html?mgpage=project&project=mgp18485). MAGs, interactive datasets, and R scripts for designing the mock communities are published on Zenodo (https://doi.org/10.5281/zenodo.7589193). Any further requests for data can be sent to.

## Acknowledgements

We thank Eitan Yaffe for fruitful discussions during the development of the pipeline. We also thank David Rogers for help with statistical methods, elucidating potential sources of sequencing artefacts that could disrupt our analysis, and generally highly insightful discussions.

