## Supplementary Materials for "Identifying and tracking mobile elements in evolving compost communities yields insights into the nanobiome"

#### I - Full description of the xenoseq pipeline

Xenoseq is a simple bioinformatic pipeline to find sequences that appear to be newly introduced into a community. Xenoseq wraps read trimming (fastp), assembly (megahit), read mapping (BWA), read filtering (samtools), and local alignment (blast), to detect putative evidence of horizontal transfer between communities.

The pipeline requires paired-end reads, and a single metadata file (tsv) which describes the names of your paired end samples (without read suffixes like "_R1.fq") in query-reference pairs, where for example the queries are evolved communities, and the references are their corresponding ancestral communities:

```
#SAMPLES (Query\tReference)
Horizontal1 Ancestral1
Horizontal2 Ancestral2
Horizontal3 Ancestral3
Horizontal4 Ancestral4
Vertical1   Ancestral1
Vertical2   Ancestral2
Vertical3   Ancestral3
Vertical4   Ancestral4
```

By default, xenoseq will look for sequences matching the names in the metadata in the directory `$PWD/samples/reads`, and will look for samples ending in `_R1.fq` and `_R2.fq`. The path to the reads and the read suffixes can be changes with the option `-p` and `-r` respectively. In other words; if you have your own reads you may need to modify the metadata and/or modify the read path/prefix options. The default output-directory is called "Xenoseq_default", and can be changed with `-o`.

Once the pipeline has parsed the metadata and verified the reads are in place, it will create the output directory ("*Xenoseq_default*"), and a subdirectory for each sample within that directory ("*Horizontal1*", "*Ancestral1*", "*Horizontal2*", etc.). For each sample in the QUERY column, it will generate the following files:

- *unique_contigs.fasta*; sequences in query not present in the reference
- *unique_contig_links_to_<REF>.tbl*; table of all contigs and the community they can be linked to (*i.e.* data for a directed graph as shown in **Figure 2b** in the main text). Requires option '-l'
- *xenotypic_contigs.fasta*; the subset of unique contigs that can be linked to another reference. Requires option '-l'
- *xenotypic_coverage*.txt*; plain text files describing the coverage of the xenotypic sequences in all other samples. Requires option '-t'
- *source_contigs*; directory with fasta files from reference samples that are themselves / are linked to MGEs

**Subroutine 0 - xenoseq_prep**

The first subroutine of xenoseq is called everytime a sample is first called by any other subroutine of the pipeline. It will use fastp to trim the reads, remove duplicate reads, and merge read pairs into a file "*merged_reads.fasta*".

**Subroutine 1 - xenoseq_find**

First, all *reference* samples are trimmed and assembled into contigs using megahit with default options. A blast database and a BWA index are prepared for each of these reference samples.

Next, for each *query*, reads are mapped against the corresponding reference in the metadata file using BWA mem, and unmapped reads are extracted using samtools (-f 4). The unmapped reads are then assembled into a file called "*unique_contigs.fasta*", which is stored in a subdirectory named after the query sample.

**Subroutine 2 - xenoseq_link**

Unique contigs for each query sample are blasted against a local database of all *other* reference samples. By default, unique contigs with at least 300 basepairs with 99% nucleotide identity are stored in "*unique_contig_links_to_<REF>.tbl*", and stored in "xenotypic_contigs.fasta". The minimal alignment length and percent identity can be modified with the `-L` and `-P` option, respectively.

**Subroutine 3 - xenoseq_trace**

Xenotypic contigs of each query sample are compared against every other sample, including the other reference samples. Coverage statistics are extracted with samtools coverage, which includes the average depth, breadth, and mapping quality.

**Running xenoseq with example data:**

After cloning the repository (and installing dependencies using the provided conda environment), the example data can be run with:

```
> ./xenoseq -m example_metadata.tsv -o Xenoseq_example -l -t
```

This example will use the reads found in samples/reads to search for xenotypic contigs in simulated data from one of our mock communities (see Methods). Unique/xenotypic contigs will then be stored in Xenoseq_example/<SAMPLE_NAME>. This example calls all the subroutines by default, but ommiting `-l` or `-t` will disable xenoseq_link or xenoseq_trace respectively.

The pipeline was written to not run any of the subroutines twice for the same sample. After running the raw pipeline (without `-l` and `-t`), rerunning it with `-l` and `-t` and the same output directory will automatically reuse previously generated files and link or trace your identified contigs.

If you want to modify any of the options (e.g. filtering thresholds, quality trimming), you can modify the relevant subroutines of xenoseq given in *xenoseq_bin/functions.sh*. Generally, we found that changing these options is not required, as further filtering can be done by analysing the coverage statistics generated by xenoseq_trace.

#### Full list of options

For an up-to-date overview of the latest pipeline and its options, see the online repository at github.com/bramvandijk88/xenoseq. Below is an overview of the options in the published version 1.0.0.

```
> xenoseq -h

----------------------------------------------------------------------
                          (XENOSEQ                                                                          v1.2.0)
----------------------------------------------------------------------

xenoseq v1.2.0

contact:

Usage:
         xenoseq -m <meta_data_tsv> -o <output_dir> -c <num_cores> -l -t

Mandatory:
        -m/--metadata                 File containing the metadata (tsv file with query-reference sets)
Optional:
        -p/--path_to_reads <STRING>       Path to reads for samples in metadata (default = samples/reads)
        -r/--read_suffix <STRING>      Read suffix corresponding to metadata n ames (e.g. when read filenames
are      Sample1_R1.fq      and      Sample1_R2.fq,      use      _R*.fq)      (default      =      _R*.fq)
        -l/--link                          After detecting unique contigs, attempt to link them to other
reference                                                                                          samples.
        -t/--trace                        After detecting xenotypic contigs, trace them across all samples.
        -c/--cores <INT>                Number of CPUs to use for smaller tasks (passed on to bwa, samtools,
etc.)                           (default                              =                              4)
        -C/--assembly_cores <INT>                Number of CPUs to use for assembly (megahit) (default = 4)
        -j/--jobs    <INT>                            Maximum  number  of  parallel  jobs  (default  =  4)
        -J/--max_assembly_jobs   <INT>     Maximum  number  of  parallel  jobs  for  assembly  (default  =  )
        -o/--output   <STRING>                                 Output   directory   to   put   all   the   data
        -L/--alignment_length           Minimal alignment length to link unique sequences to other reference
samples.
        -S/--single_end <STRING>         Assume single-end reads (e.g. use only Sample1_R1.fq and skip read
merging)
        -P/--alignment_pid              Minimal percent identity to link unique sequences to other reference
samples.
        -f/--force_relink                Link unique sequences to reference samples, even when this step is
already performed.
```

### II - Benchmarking ‘xenoseq_find’

To benchmark the xenoseq_find subroutine, we simulated the introduction of MGEs into mock-communities. For this, six known genomes were downloaded from RefSeq (*Azospirillum thiophilum*, NZ_CP012401.1; *Burkholderia pseudomallei*, NC_006350.1; *Cytophaga hutchinsonii*, NC_008255.1; *Paenibacillus donghaensis*, NZ_CP021780.1; *Pseudomonas fluorescens SBW25*, AM181176.4; and *Rhodanobacter denitrificans*, NC_020541.1), as well as a small collection of plasmid and phage genomes. Then, two types mock-communities were generated, one with an even taxon distribution (**Figure S2.1, left-hand side**), and one with a highly skewed taxon distribution (**Figure S2.1, right-hand side**). Note that the taxon distribution in compost communities as discussed in the main text is much more skewed, and may therefore be an even harder dataset.

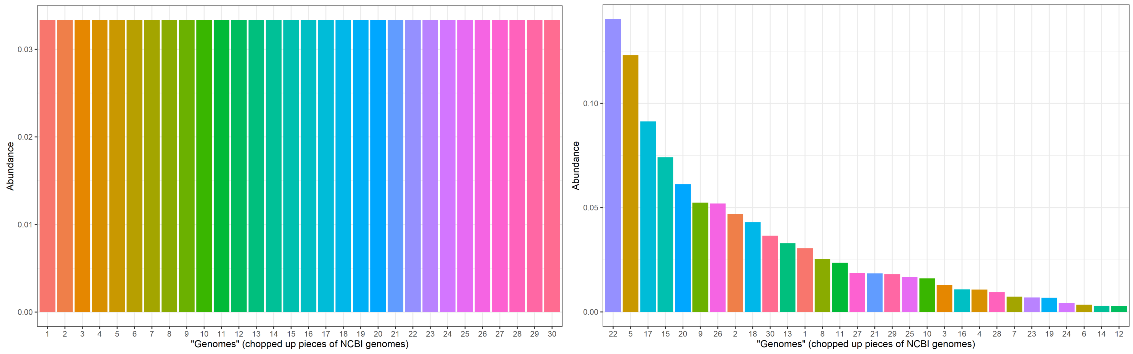


*Figure S2.1 Mock community taxon distributions (left: even dataset, right: skewed dataset)*

For computational efficiency, the taxa in the mock communities were not whole genomes, but chopped up pieces of chromosomal DNA of, on average, 80kb long (**Figure S2.2**)

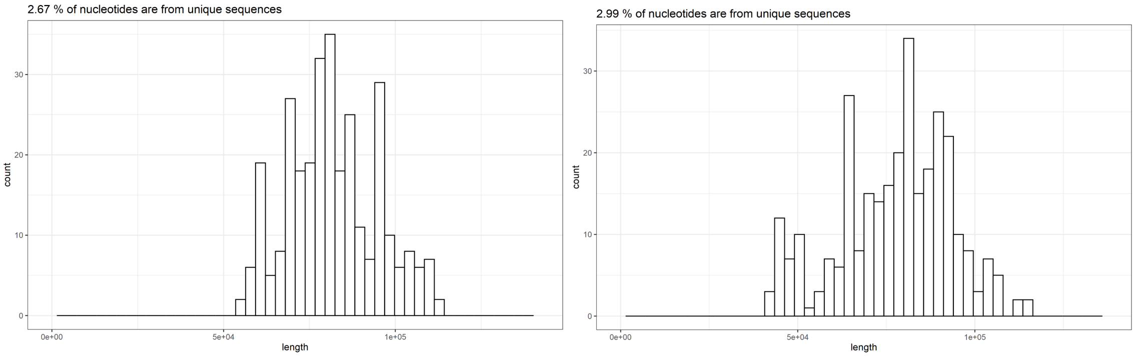


*Figure S2.2 Mock community genome size distributions (left: even dataset, right: skewed dataset)*

Next, MGE sequences were either i) randomly inserted into the genomes (shown in red below), or ii) included as separate replicons linked to a single genome (shown in green below). Some generations of neutral evolution are simulated to allow certain MGEs to amplify, whereas others may get lost again. The final distribution of MGEs in the mock communities is shown in **Figure S2.3**.

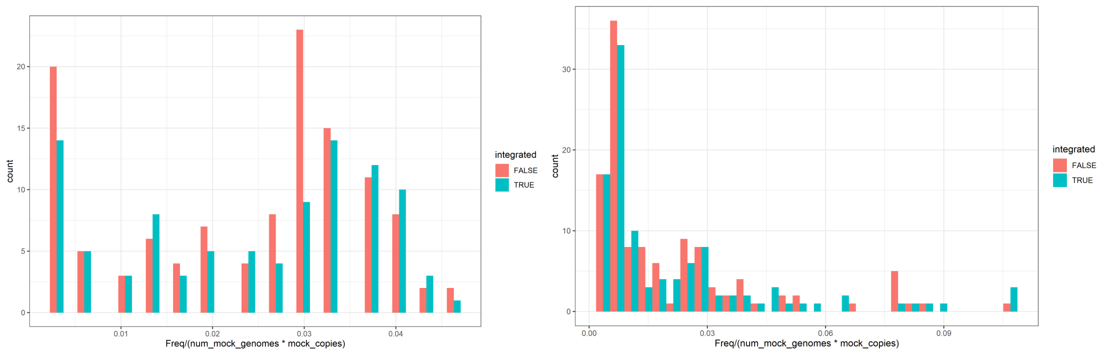


***Figure S2.3** (left: MGE abundance in even dataset, right: MGE abundance in skewed dataset)*

Illumina reads were generated from the resulting genomes/MGEs, which were used as input to benchmark the pipeline. Simulations of mock-genomes as described above was done in R using the packages biostrings and seqinr. Artificial illumina reads were generated with ART, using both default and ten-fold elevated error rates.

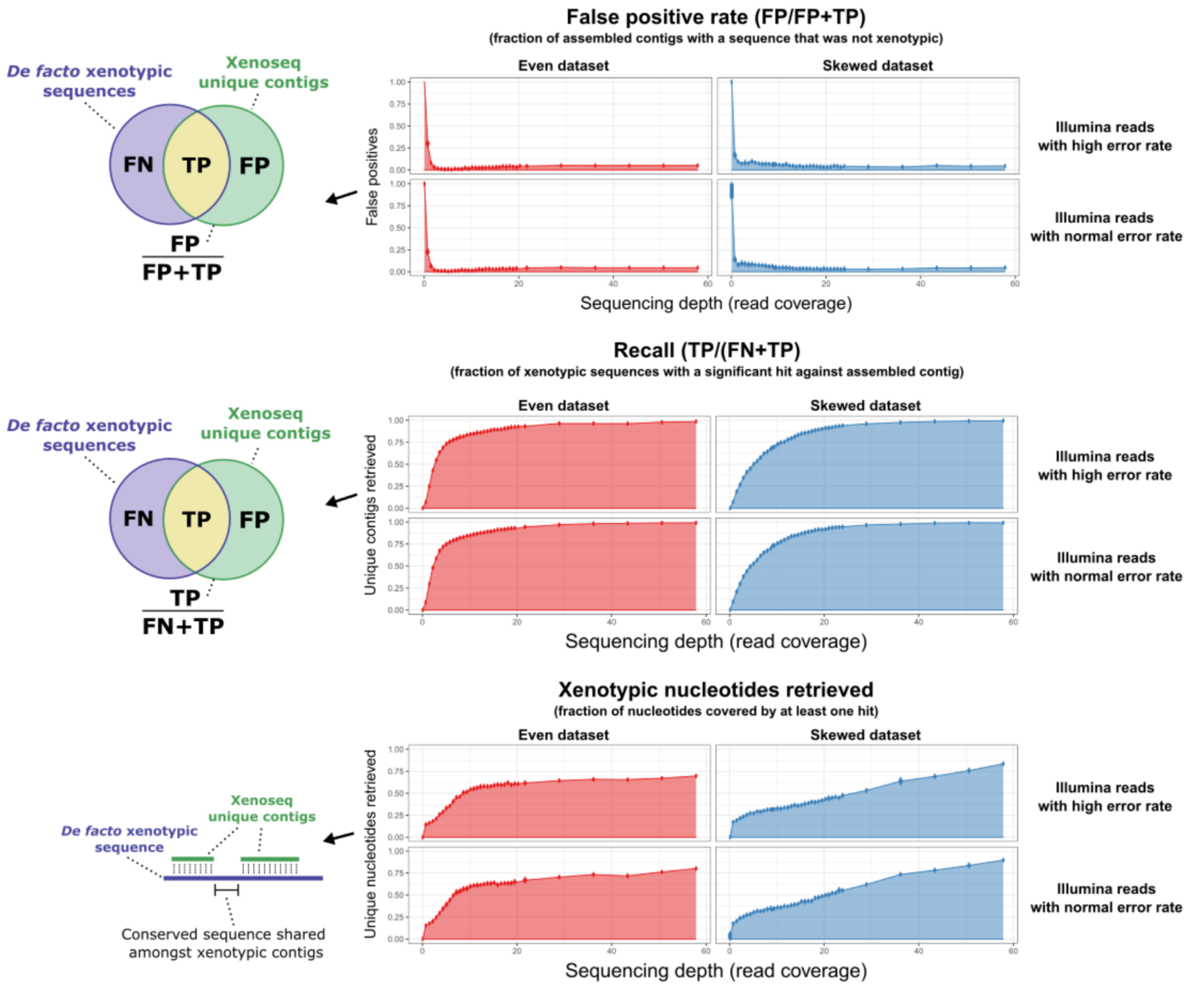


**Figure S2.4:** Benchmark of false positive rate, recall, and fraction of retrieved nucleotides using the xenoseq on mock community data. For both mock community datasets (even and skewed), Illumina reads were generated using ART . Using a range of sequencing depths (-f option, x-axis), error rates (-q option, -3 for top panels, 0 for bottom panels), and random number seeds (-s option ranging from 1 to 5). These simulated Illumina reads where used as input for the Xenoseq pipeline to calculate false positive rates (top panel), recall values (middle panel), and the fraction of retrieved nucleotides (bottom panel).

The results of the benchmarks are shown in the Figure S2.4. We found that our pipeline has low false positive rates, even at minimal coverage. Only when coverage is 1 or lower, does the false discovery rate increase. Interestingly, the false positive rate does not converge to zero even at high coverage. We found that this is explained by reads that are not mapped by BWA, but nevertheless assembled using Megahit. It is important to note that, although the false positive rate is fairly low in all test cases, it is likely that many biological samples contain many sequences that will be poorly covered. Therefore, identifying a sequence as “unique” using the xenoseq pipeline is insufficient reason to assume transfer. The xenoseq_link subroutine (described above) largely circumvents this by only retaining sequences that are highly similar to sequences in other reference communities. The false positive rate can be further decreased by increasing the threshold for linking sequences to ancestral communities (options -L and -P).

Next, the recall rate shows what proportion of true negatives is retrieved. While recall increases quite rapidly with increasing coverage, it only approaches unity at high coverage, suggesting that some sequences can be missed. We found that this is due to MGEs with similar structure present in the reference set, which masks the introduction of a new sequence into the community even at high coverage. A similar problem may arise when MGEs share homologous regions, as “unique contigs” are likely to be split at these positions. It is therefore expected that xenoseq will return any biologically interesting sequences in parts, and not as a closed contig. We found that, when the unique contig is an independent replicon (i.e. not integrated into the chromosome, like a phage or plasmid), this sequence is likely found in full in the corresponding reference community assembly. However, if the MGE was integrated into a genome, one needs to align the sequence to fully investigate its origin.

Taken together, the benchmark revealed that the false positive rate is generally acceptable, although sequences in biological communities are likely to hide under the metagenomic detection level. Therefore, we developed extra subroutines to ensure that contigs are of xenotypic origin (see Methods and Supplementary Material I)

### III – Additional supplementary figures

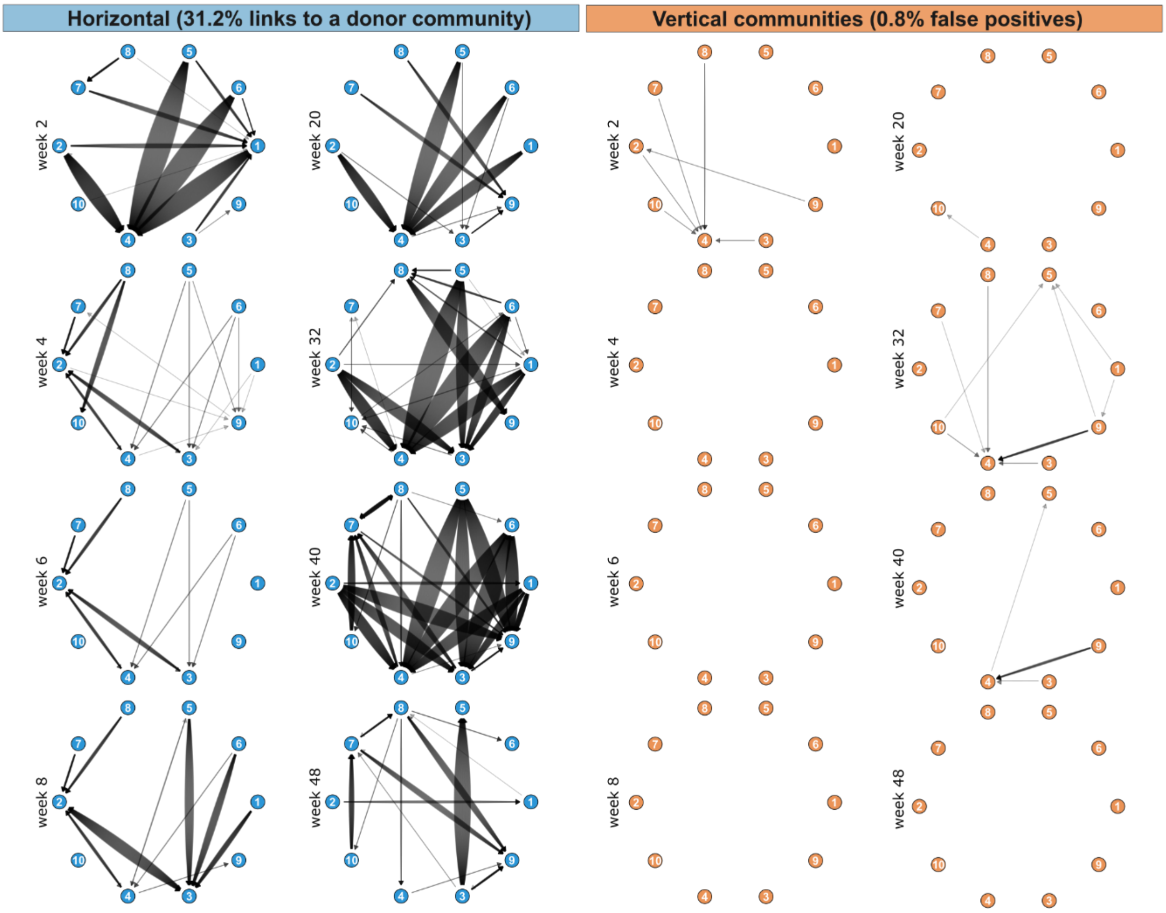


**Supplementary Figure 1: Many more xenotypic sequences are inferred from horizontal communities than from vertical communities.** On the left-hand side, unique sequences from all samples from Quistad et al., (2020) are linked to ancestral communities. Every arrow is a blast hit of at least 300 nucleotides long with at least 99% nucleotide identity. On the right-hand side, the same is shown for vertical communities. The arrows remaining in the vertical communities are, by virtue of the experimental design, not xenotypic, and should be regarded as false positives.

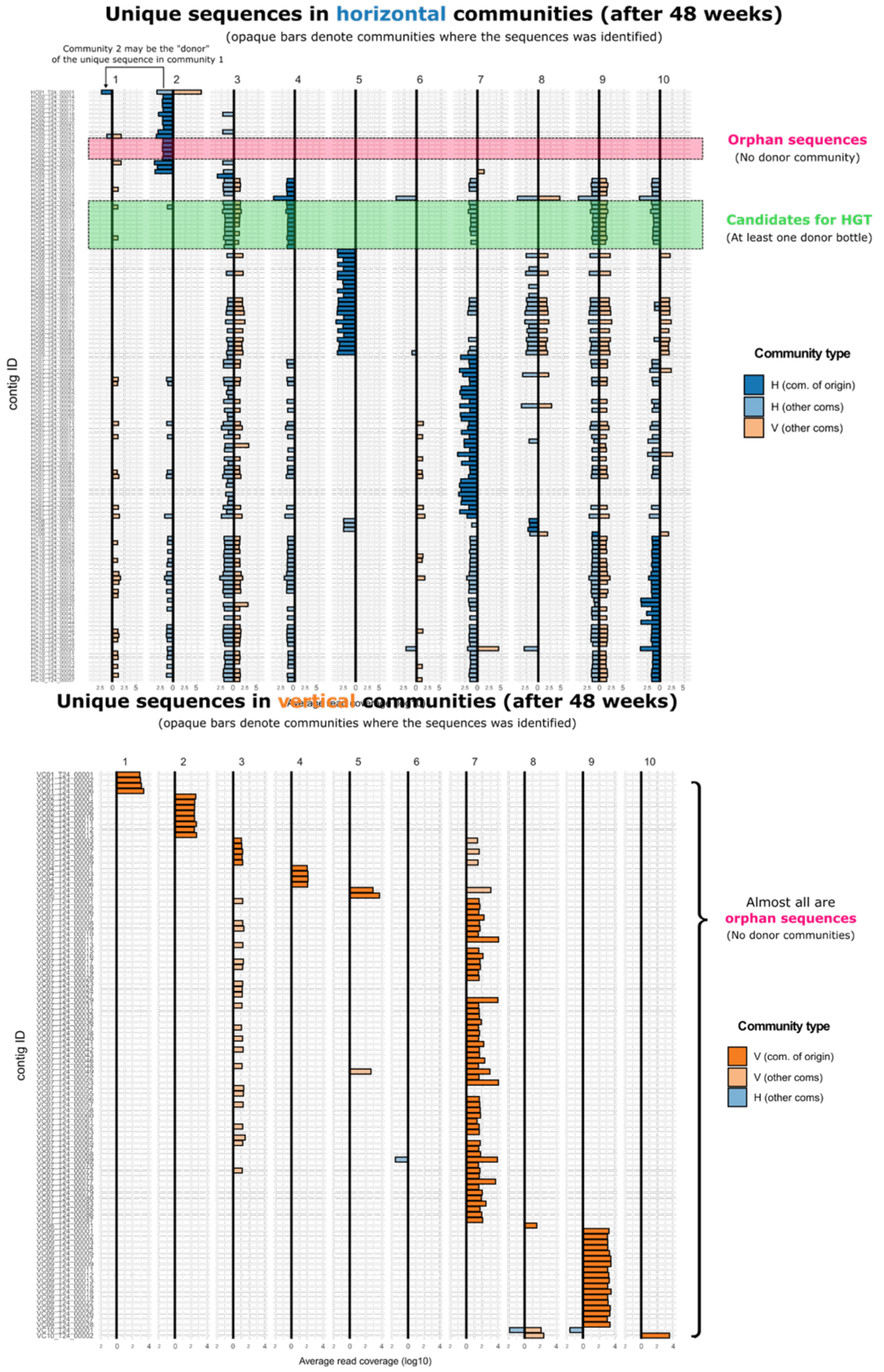


**Supplementary Figure 2: Linking unique sequences from 'xenoseq_find' to ancestral communities by read mapping** Similar to Supplementary Figure 1, but using read mapping instead of blastn to visualise the links shown in **Supplementary Figure 1** and **Figure 2b**. In the top panel, unique sequences from horizontal communities are mapped against other ancestral communities. Many sequences can be linked to at least one "donor" community. In the bottom panel, unique sequences from vertical communities are mapped against other ancestral communities, showing that most of them are "orphans".

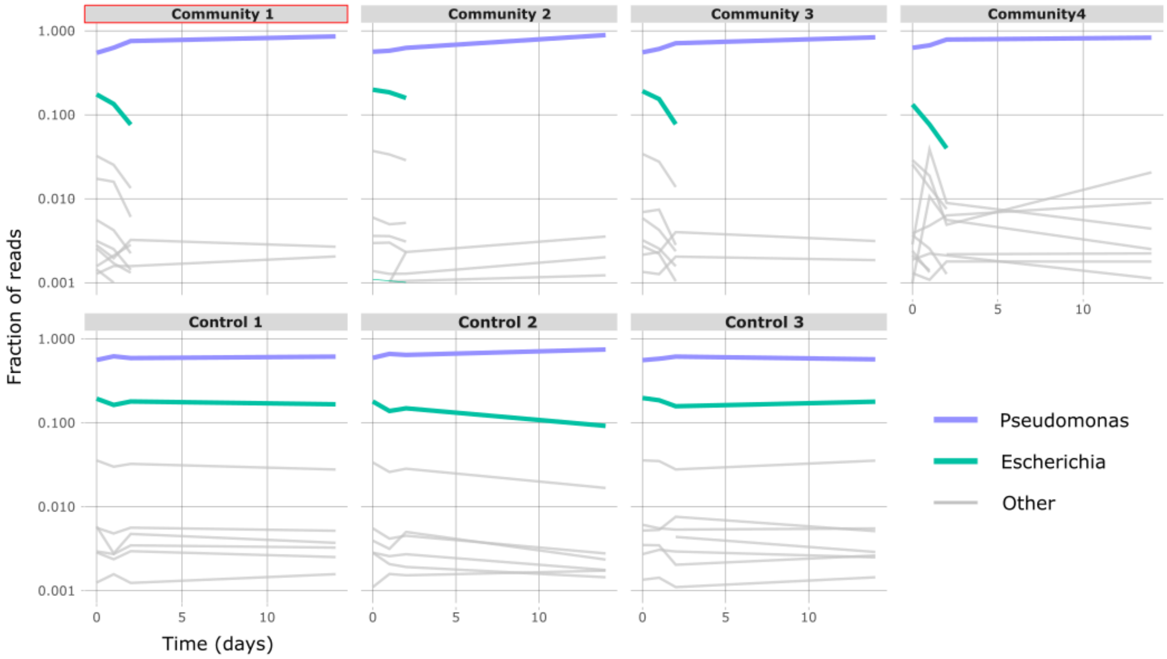


**Supplementary Figure 3: Free DNA is rapidly degraded in the presence of compost communities.** Four compost communities and three controls (see Methods) were spiked with high concentrations of *Escherichia coli* DNA at day 0, and spiked with *Pseudomonas fluorescens* SBW25 DNA daily. Reads were annotated with Diamond using the non-redundant (NR)[65] protein database. The data indicate that *E. coli* DNA was rapidly degraded, but only in the presence of compost communities (top row). When no community was present (bottom row), the *E. coli* DNA that was spiked at day 0 remained unchanged over time.

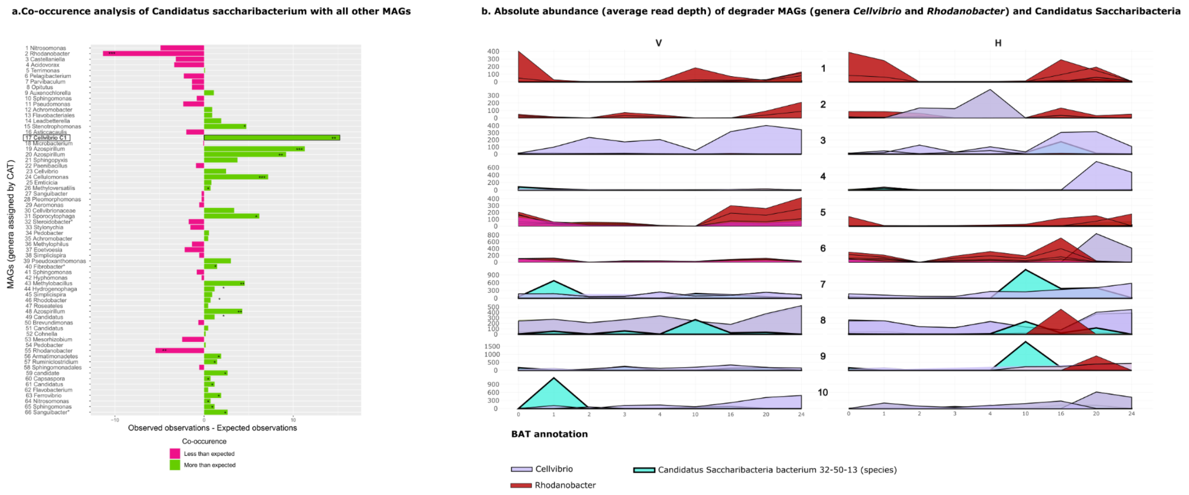


**Supplementary Figure 4: MAG annotated as *Candidatus saccharibacterium* is most likely associated with Cellvibrio C1, and shows massive blooms in the community, suggesting a parasitic lifestyle.**

**a.** For various MAGs, it was tested how often it co-occurs with *Candidatus saccharibacterium* within a sample (observed co-occurences) , and how often one would one expect to see co-occurrence given the number of observations across all samples (i.e., the expected co-occurrence based on chance). The difference between these two numbers is plotted on the x-axis, shown in purple when co-occurrence is less frequent than expected by chance, and green when co-occurrence is more frequent than expected by chance. For each observation on the x-axis, p-values represent the probability of finding at least that many observations given the frequencies of the respective MAG and *C. saccharibacterium* (* = p-value < 0.05, ** = p-value < 0.01, *** = p-value < 0.001). **b.** The absolute read coverage is shown for relevant degrader genera (*Cellvibrio* and *Rhodanobacter*) and *Candidatus saccharibacterium*. This reveals that *C. saccharibacteriu*m shows massive blooms in communities which have *Cellvibrio spp.* as the primary degrader. In community 8 and 9, a bloom of *C. saccharibacterium* briefly causes *Rhodanobacter* to dominate.

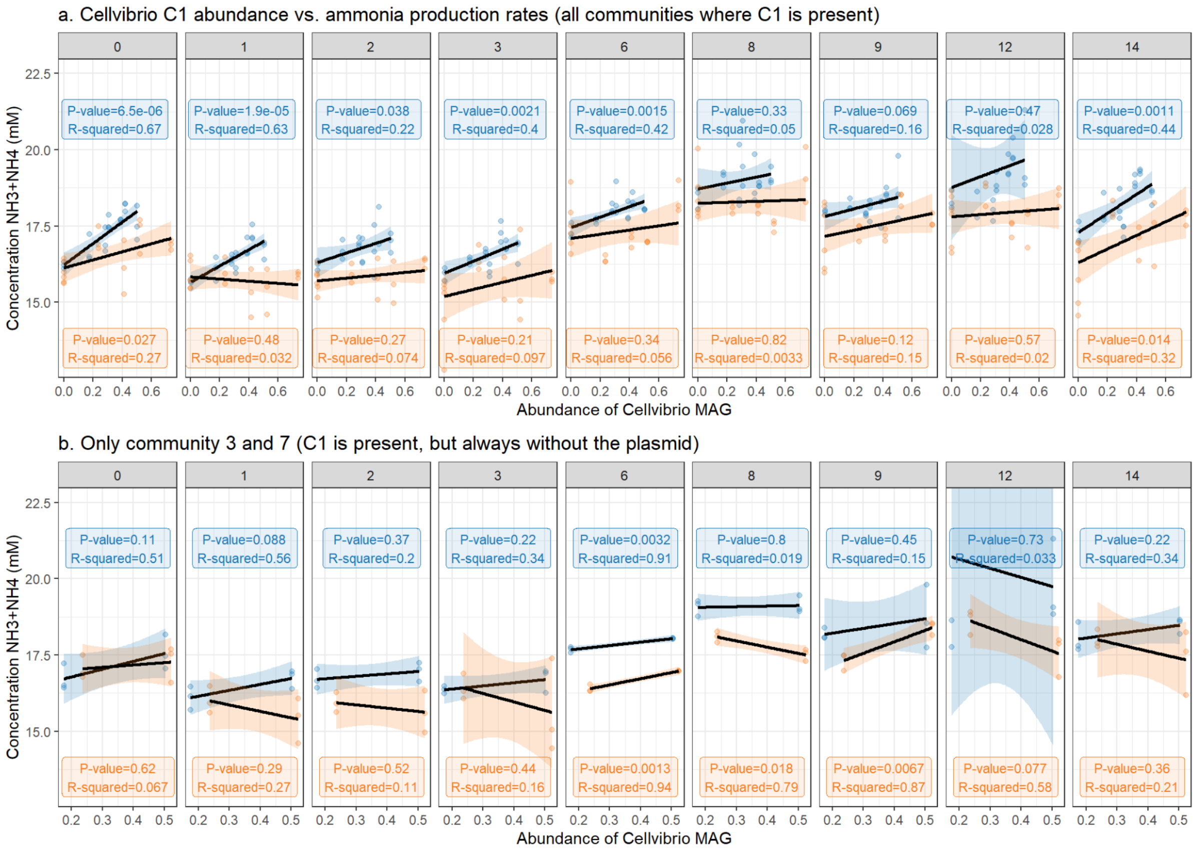


**Supplementary Figure 5 - Communities abundant in nitrogen-fixing *Cellvibrio* MAGs (C1) carrying the CP plasmid-like element are enriched in ammonium, especially shortly after transfer to fresh media.** The relative abundance of *Cellvibrio* C1 was determined by the proportion of reads mapping against this MAG after 48 weeks, and plotted against ammonia production measurements from Quistad *et al.* (2020). *Cellvibrio* C1 was present in four vertical communities and seven horizontal communities, and plotted on the x-axis. Note that these measures on the x-axis are identical for each day, as it was obtained from a single metagenomic sample. On the y-axis, ammonia concentrations (NH3+NH4) are plotted, with three technical replicates for each community per day, and a linear regression model was fitted to the data points. In panel b, only communities are shown where the plasmid was not present, greatly affecting the pattern observed in panel a. Taken together, this suggests that the Cp plasmid-like element promotes nitrogen fixation in *Cellvibrio* C1.

#### IV – Online supplementary data

##### Published metagenomic data

All MAGs, including separate assemblies of *Candidatus saccharibacteria*, *Cellvibrio spp.*, and the identified Cp plasmid-like element are uploaded to zenodo.

##### Supplementary table I

Supplementary Table I consists of an Microsoft Excel document with four Work sheets. Work sheet I summarises the number of identified unique contigs and xenotypic contigs across all analysed compost samples. Work sheet II gives an overview of all xenotypic contigs, including their length, kmer coverage, in which community they were identified, and what taxon was assigned by Contig Annotation Tool (CAT). Work sheet III shows the data used for **Figure 3c**, where xenotypic sequences are investigated for enrichment of MGEs by comparing them with arbitrary sequences from the whole community. Work sheet IV shows measurements performed on MGE cocktails. Finally, Work sheet V gives the BAT annotation for all MAGs as shown in **Figure 5a-b**, the estimated completion and contamination (CheckM), and the assigned metabolic function scores (cellulose degrader, cellulose scavenger, nitrogen fixer (I), nitrogen fixer (II), nitrate reducer, and nitrite reducer).
